# Genomic and phenotypic characterization of experimentally selected resistant *Leishmania donovani* reveals a role for dynamin-1 like protein in the mechanism of resistance to a novel anti-leishmanial compound

**DOI:** 10.1101/2021.01.05.425522

**Authors:** Aya Hefnawy, Gabriel Negreira, Marlene Jara, James A. Cotton, Ilse Maes, Erika D’ Haenens, Hideo Imamura, Bart Cuypers, Pieter Monsieurs, Christina Mouchtoglou, Hans De Winter, Matt Berriman, Mandy Sanders, Julio Martin, Geraldine de Muylder, Jean-Claude Dujardin, Yann G.-J. Sterckx, Malgorzata Anna Domagalska

## Abstract

The implementation of prospective drug resistance (DR) studies in the R&D pipelines is a common practice for many infectious diseases, but not for Neglected Tropical Diseases. Here, we explored and demonstrated the importance of this approach, using as paradigms *Leishmania donovani*, the etiological agent of Visceral Leishmaniasis (VL), and TCMDC-143345, a promising compound of the GSK ‘Leishbox’ to treat VL. We experimentally selected resistance to TCMDC-143345 in vitro and characterized resistant parasites at genomic and phenotypic levels. We found that it took more time to develop resistance to TCMDC-143345 than to other drugs in clinical use and that there was no cross resistance to these drugs, suggesting a new and unique mechanism. By whole genome sequencing, we found two mutations in the gene encoding the *L. donovani* dynamin-1-like protein (LdoDLP1) that were fixed at highest drug pressure. Through phylogenetic analysis, we identified LdoDLP1 as a family member of the dynamin-related proteins, a group of proteins that impacts the shapes of biological membranes by mediating fusion and fission events, with a putative role in mitochondrial fission. We found that *L. donovani* lines genetically engineered to harbor the two identified LdoDLP1 mutations were resistant to TCMDC-143345 and displayed altered mitochondrial properties. By homology modeling, we showed how the two LdoDLP1 mutations may influence protein structure and function. Taken together, our data reveal a clear involvement of LdoDLP1 in the adaptation/resistance of *L. donovani* to TCMDC-143345.

**Importance:** Humans and their pathogens are continuously locked in a molecular arms race during which the eventual emergence of pathogen drug resistance (DR) seems inevitable. For neglected tropical diseases (NTDs), DR is generally studied retrospectively, once it has already been established in clinical settings. We previously recommended to keep one step ahead in the host-pathogen arms race and implement prospective DR studies in the R&D pipeline, a common practice for many infectious diseases, but not for NTDs. Here, using *Leishmania donovani*, the etiological agent of Visceral Leishmaniasis (VL), and TCMDC-143345, a promising compound of the GSK ‘Leishbox’ to treat VL, as paradigms, we experimentally selected resistance to the compound and proceeded to genomic and phenotypic characterization of DR parasites. The results gathered in the present study suggest a new DR mechanism involving the *L. donovani* dynamin-1 like protein (LdoDLP1) and demonstrate the practical relevance of prospective DR studies.

## Introduction

The lifespan of any anti-microbial drug is, unfortunately, limited; its clinical use generally represents a new step in the arms race between human creativity and pathogen adaptability. Sooner or later, drug resistance (DR) or another phenotypic adaptation arises (1). Human countermeasures (such as new therapeutic regimens or combination therapy) can be adopted, but the drug will ultimately have to be replaced by a new compound, if any are available. Understanding the process of DR and developing strategies to counter it are particularly critical for neglected tropical diseases (NTDs), for which there are typically only a few drugs in the therapeutic arsenal and the R&D pipeline (2). It is essential to safeguard existing compounds and to develop new ones. At the same time, studies on molecular mechanisms of resistance are classically applied to understand the mode of action of antimicrobial agents, because of the possibility that the resistance determinants are caused by specific genetic variations resulting in the altered target binding to the drug molecule.

For NTDs, DR is generally studied retrospectively, once DR has already been established in clinical settings. In a recent opinion paper, we recommended to keep one step ahead in the arms race and implement prospective DR studies in the R&D pipeline, a common practice for many infectious diseases, but not for NTDs. Two specific recommendations have been given so far (2): (i) exploiting resources of parasite bio-banks to test the efficacy of novel compounds on a wider range of parasites, including recent isolates from clinically relevant settings for the prospective use of the compound -a practice shown to be highly relevant (3)- and (ii) experimentally selecting DR to new lead compounds and characterizing it broadly to assess the adaptive skills of the parasite to the compound, guide further drug development and help countering DR if it develops in clinical practice.

Here, using *L. donovani* (the etiological agent of Visceral Leishmaniasis, VL, which is fatal if left untreated) and TCMDC-143345, a promising anti-VL compound of the GSK ‘Leishbox’ (4) as paradigms, we experimentally selected resistance to the compound and proceeded to a genomic and phenotypic characterization of DR parasites. We found that it took more time to develop resistance to TCMDC-143345 than to other drugs in clinical use and that there was no cross resistance to these drugs, suggesting a new and unique mechanism. Whole genome characterization of independent TCMDC-143345-resistant lines highlighted two mutations in the gene encoding dynamin-1-like protein (LdoDLP1) that were fixed at highest drug pressure. Through phylogenetic analysis, we identified LdoDLP1 as a family member of the dynamin-related proteins (DRPs), a group of proteins that impacts the shapes of biological membranes by mediating fusion and fission events, with a putative role in mitochondrial fission. We genetically engineered our *L. donovani* strain to harbor the two identified LdoDLP1 mutations: parasites were resistant to TCMDC-143345 and displayed altered mitochondrial properties. The results are further supported by homology modeling which provides insights as to how the two LdoDLP1 mutations may influence protein structure and function. Taken together, the data presented in this paper reveal a clear involvement of LdoDLP1 in the adaptation/resistance of *L. donovani* to TCMDC-143345. Our results also demonstrate the practical relevance of prospective drug resistance studies to guide R&D pipeline and future clinical applications of that compound.

## Materials and Methods

### Parasites

We used the *L. donovani* strain MHOM/NP/03/BPK282/0 clone 4 (further called LdBPK_282 cl4, see growth conditions in suppl. text), originally derived from a Nepalese patient with confirmed VL and cryo-preserved at the Institute of Tropical Medicine, in Antwerp, Belgium. The strain is considered sensitive to antimonials [Sb^III^], Miltefosine [MIL] and Amphotericin B [AmphoB] and it was used for determining the reference genome of *L. donovani* (5). The strain kept most of its intrinsic phenotypic features, like virulence (5), transmissibility to sand flies (6) and natural drug susceptibility (7), thus constituting a good model for ‘real-life’ parasites that will be exposed to new drugs in natural conditions.

### Selection of resistance and stability of resistance

For the selection of resistance to TCMDC-143345, promastigotes were initially grown in quadruplicates (lines A, B, C, D) and later on because loss of lines A and B, line D was divided in 4, constituting a total of 5 lines (C, D1-D4). MIL was used as positive control for the experimental set-up of drug resistance selection and two duplicates were used (A and B). As negative controls, two additional lines were used: (i) the Wild-Type [WT] line LdBPK_282 cl4 maintained during the same passage numbers as the resistant lines but without the drug pressure and (ii) a WT line maintained with DMSO (which was used as solvent for TCMDC-143345). The resistant lines were maintained in the continuous presence of drugs, as described elsewhere (7). Increasing concentrations of drugs were added in a step-wise manner until all lines grew at similar rates as wild-type parasites: (i) for MIL: 0, 2, 10, 15, 60, 100 μM, (ii) for TCMDC-143345, line C: 0, 0.2, 1, 2, 4, 5, 6, 8, 12 and 12 μM and for line D: 0, 0.2, 1, 2, 4, 5, 6, 10, 18 and 25 μM. Each selection round was approximately 5 weeks (2 passages per week) with the IC_50_ measured after each round. The selection flowchart is summarized in figure S1. To test the stability of the TCMDC-143345-resistant phenotype, the resistant line D1 was maintained for 20 weeks without drug pressure, after which the IC_50_ was measured.

### Promastigotes susceptibility tests

Susceptibility tests were performed after each selection round (drug resistance selection) or after parasite engineering (CRISPR-Cas9 or over-expression, see below). The IC_50s_ were determined on logarithmic-stage promastigotes after 72 hours of exposure to TCMDC-143345 or MIL with a resazurin assay as previously described (8) and summarized in Supplementary text. For the cross-resistance experiments the same protocol was used. The following maximal concentrations were used for the testing of the compounds: 50 µM for TCMDC-143345 and compound Y, 400 μM for MIL, 2 mM for Sb^III^ and 200 μM for AmphoB. Ten points of 1 in 2 dilutions were used per compound. Four independent experiments were run with technical duplicates per experiment.

### Amastigotes susceptibility tests

Phorbol myristate acetate (30 nM) (PMA, Sigma) was added to THP-1 cells (human monocytic leukemia, ATCC-TIB-202, see maintenance conditions in supp. text) at 37°C for 48 hours to differentiate these into adherent macrophages. Cells were washed and incubated with complete RPMI medium containing stationary phase (day 6) *L. donovani* promastigotes at a macrophage/promastigote ratio of 1/30. After 5 h incubation at 37°C, non-internalized promastigotes were removed by 3 successive washes with PBS and infected macrophages were incubated with TCMDC-143345 in RPMI medium supplemented by 5% heat inactivated Horse serum for 96 h. TCMDC-143345 was tested with a starting concentration of 25 μM in a 3-fold serial dilution. A 3-fold serial dilution of 3 μM amphotericin B was used as positive control. Experiments were done in triplicate with technical duplicates per experiment. For confocal microscopy, infected cells were washed with PBS, fixed for 30 minutes with 4% formaldehyde, rinsed again with PBS and stained with 4’,6’-diamidino-2-phenylindole (DAPI 300 nM). Images were acquired with an LSM 700 Zeiss confocal microscope. The number of infected macrophages and the number of amastigotes per infected macrophage were determined by manual counting. These numbers obtained from the average of counted wells were used to establish the infection index (% infected macrophages × amastigotes/infected macrophages). IC_50s_ were calculated with GraphPad Prism using a sigmoidal dose-response model with variable slope.

### DNA and library preparation for whole genome sequencing

At each round of the resistance selection (5 weeks culture), parasites were harvested from lines C and D and from the two WT controls (maintained without and with DMSO). The list of samples sequenced is summarized in Table S1. DNA isolation was done using QIAamp DNA blood minikit (Qiagen), and the DNA concentration was assessed with the Qubit DNA broad-range DNA quantification kit (Thermo Fisher). Library preparation and sequencing of the different lines of the stepwise selection were performed at the Wellcome Sanger Institute (Hinxton, United Kingdom). Genomic DNA was sheared into 400–600-base pair fragments by focused ultrasonication (Covaris Adaptive Focused Acoustics technology, AFA Inc., Woburn, USA). Amplification-free indexed Illumina libraries were prepared (9) using the NEBNext Ultra II DNA Library Prep kit (New England BioLabs). The libraries were quantified using the Accuclear Ultra High Sensitivity dsDNA Quantitative kit (Biotium) and then pooled in equimolar amounts. Paired end reads of 150-bp were generated on the Illumina HiSeq X10 according to the manufacturer’s standard sequencing protocol (10). Data Release: raw data was deposited in the European Nucleotide Archive (ENA) with the accession number ERS441806-ERS441816.

### Whole genome sequencing data analysis

Somy, single nucleotide polymorphisms (SNPs), local copy number variations (CNVs) and Indels were determined as described elsewhere (5),(11) using the BPK282v2 PacBio reference genome (5); more details can be found in supplementary text. SNPs and small indels were considered significantly different between parasite lines when the allele frequency showed difference of at least 0.25 and Mann–Whitney U test p-value <0.05 (12). Allele frequency shifts larger than 0.80 were considered homozygous variants. We used one criterion to evaluate whether a gene or chromosome copy number difference was biologically meaningful and statistically significant: the absolute difference in gene/chromosome copy number should be at least 0.5 to be significant. Gene ontology analyses were done as explained in supplementary text. Heat maps were created using the heatmap3 package in R (R Development Core Team 2015). Reasoning that mutations away from the sensitive parental strain at causative loci were likely to contribute to resistance but that multiple variants in the same loci could contribute to the resistance phenotype, we adopted a simple approach to identify significant loci informed by burden tests used in rare-variant association studies (13). We summed the non-reference allele frequency of variants at each locus for each sequence sample and tested for association by regression of these total allele frequencies per locus against the measured IC_50_ for that sample.

### CRISPR-Cas9 mediated engineering of Leishmania

LdBPK_282 cl4 parasites were transfected with the linearized pTB007 vector, obtained from Dr. Eva Gluenz (University of Oxford, UK) (14). Transgenic parasites were selected with 25µg/mL Hygromycin starting 24h after transfection and a clone was isolated using a micro-drop method (15). In order to introduce the Ala324Thr or the Glu655Asp mutation using the CRISPR-Cas9 system, the single guide RNAs (sgRNAs) DynMut1-gRNA and DynMut2-gRNA were designed targeting the sequences CAGCAGCTGTGC<u>A</u>GTGGGCT and GGCACTGCTCTCCGA<u>G</u>CCCCC respectively (sites of mutation are underlined). The sgRNA templates for in vivo transcription were generated by PCR as previously described (14). The sequence of all primers used in this work are provided in Table S2. For each mutation, a double-stranded donor DNA bearing the missense mutation was generated by annealing of synthetic oligos. The DNA repair templates also included synonymous nucleotide substitutions to distinguish the CRISPR-Cas9-mediated mutations from potential naturally occurring mutations. The Ala324Thr and the Glu655Asp mutations were independently recreated by transfecting the respective sgRNA template and donor dsDNA using the Basic Parasite Nucleofector™ Kit 1 (Lonza) with the U-033 program following manufacturer recommendations. Control transfections were made by transfecting either each donor dsDNA without their respective sgRNA templates or by replacing each donor dsDNA by the DynWT1 or DynWT2 dsDNAs, which lack the missense mutation. After 24h post transfection, 10^6^ parasites of each transfection were transferred to a 24-wells plate in a final volume of 1mL/well of HOMEM medium with 20% Fetal Bovine Serum and 9µM of TCMDC-143345 or 0,1% DMSO (control). Plates were incubated at 26°C for 12 days and cell viability was determined by flow cytometry using the NucRed™ Dead 647 ReadyProbes™ and the Vybrant™ DyeCycle™ Green dyes (ThermoFisher). Parasites that survived and grew in the presence of TCMDC-143345 were transferred to culture flasks and kept under pressure with 6µM of TCMDC-143345 for 2 passages, when clones were isolated with a microdrop method (15) and grown in absence of drug pressure.

### Over-expression of WT LdoDLP1 gene in Lines C and D3

The wild-type LdoDLP1 gene was PCR-amplified from the gDNA of LdBPK_282 cl4 with the primers InF-LdDNM1-F and InF-LdDNM1-R and cloned in the *Not*I and *Nco*I sites of the pLEXSy-Hyg2.1 expression vector (Jena Bioscience). The plasmid was linearized with the *Swa*I enzyme and transfected in parasites of fully resistant Lines C and D3, as well as the standard LdBPK_282 cl4 using the Basic Parasite Nucleofector™ Kit 1 (Lonza) with the U-033 program. The empty, linearized pLEXSy-Hyg2.1 vector was also transfected in each line as control. Parasites were selected and maintained with 50µg/mL Hygromycin after 24h post-transfection.

### Phylogenetic analysis

The amino acid sequences of several DRPs (see details in supplementary text) were aligned using MAFFT (16) to generate a sequence alignment from which a rooted phylogenetic tree was constructed through the maximum-likelihood method using PHYML (17). *Escherichia coli* CrfC, which shares features with the DRP family members, was employed as an outgroup to root the phylogenetic tree. The reliability of the tree was verified by performing 1000 bootstrap replicates.

### Mitochondrial membrane potential and cell viability

[i] Selected resistant lines C and D3, together with DMSO control with same number of passages as well as [ii] CRISPR-engineered mutants, together with the WT Line and the WT transfected with the CRISPR-Cas9 and called pT007 (both used as controls) were cultivated at a density of 1 x 10^6^ without or with 12.5 μM of TCMDC-143345. On days 2, 4 and 7 the mitochondrial membrane potential (MtMP) and cell viability were co-evaluated with the Mitotracker DeepRed and NucGreen respectively (Thermo Fisher Scientific). Briefly, 1 volume of parasites was incubated with 2 volumes of a medium containing 0.1 μM of cell tracker Deep Red and 1 drop/mL of NucGreen. The samples were incubated for 15 min at 26 °C and subsequently re-pelleted by centrifugation at 1500 g per 5 min. The cells were resuspended in new medium and analysed by flow cytometry (BD FACS Verse) in the medium flow rate mode. An unstained sample was included in each experiment as negative control for the establishment of the autofluorescence and the gates for the selection of the population positive and negative for both fluorochromes.

### Homology modeling

A homology model for the LdoDLP1 dimer was generated using MODELLER (18) with the following crystal structures as templates: *Homo sapiens* DLP1 (Uniprot ID O00429, PDB ID 4BEJ) (19), *Rattus norvegicus* DNM1 (Uniprot ID P21575, PDB ID 3ZVR)(20), *H. sapiens* DNM3 (Uniprot ID Q9UQ16, PDB ID 5A3F) (21) and *H. sapiens* DNM1 (Uniprot ID Q05193, PDB ID 3SNH) (22). The homology model has zDOPE and GA341 scores of −0.255 and 1.000, respectively, thereby indicating its reliability. Molecular graphics visualization and analysis were performed with UCSF ChimeraX (23).

## Results

### Experimental resistance of promastigotes to TCMDC-143345 takes 50 weeks to establish, is stable, and is maintained in amastigotes

To assess the ability of parasites to adapt to TCMDC-143345 we set up an experimental resistance experiment in quadruplicates for TCMDC-143345 (Line A, B, C, and D), starting with 0.2 µM and for comparison we used Miltefosine (MIL) pressure in duplicates (lines A and B) starting with 2 µM. For TCMDC-143345, two lines (A and B) were lost during the selection process after round 4. The selection was continued from round 5 with line C and lines D1-D4, generated by splitting the original line D at this round (Fig. S1, Table S1).

The first parameter to be evaluated was the time to resistance (7), which was defined as the time needed for each line to display a wild-type (WT) growth curve in the presence of the highest selection pressure following the stepwise selection process. The selection dynamics for various lines are shown in Figure 1A. It took 10 selection rounds (time to resistance of approximatively 50 weeks) for line D1 to reach the highest TCMDC-143345 IC_50_ (55 µM). In comparison, the IC_50_ of TCMDC-143345 for the WT line maintained without drug pressure for the same time was 2-3 µM. The selection dynamics for line C was rather different from line D1 and the resistance selection was not complete even after 10 rounds, thereby implying a longer time to resistance. At round 5 of the selection, line C was not adapting well, which is why the pressure was decreased. The highest IC_50_ achieved was 15 µM at round 9. For the remaining D replicates, line D3 had a similar resistance profile as line D1 while for lines D2 and D4 the selection had to be stopped one round earlier (IC_50_ ranging between 16 and 25 µM, Fig. S1) due to the limited availability of TCMDC-143345.

**Fig 1.**
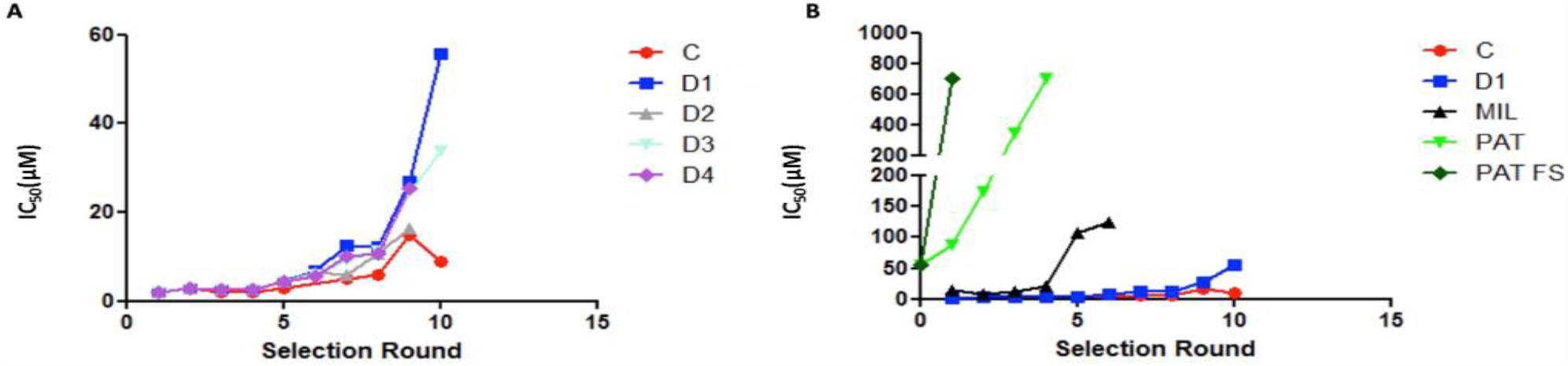
Time to resistance during selection experiments. The IC_50_ is shown in µM on the Y axis and selection round on the X axis (approx. 5 weeks per round) in (A) the different lines of LdBPK_282 cl4 kept under pressure with TCMDC-143345, in (B) line C and D1 are compared with LdBPK_282 cl4 selected for resistance to different known compounds (Miltefosine, MIL; potassium antimonyl tartrate, PAT); PAT FS, consisting in a ‘flash’ selection exposing the parasites directly to the highest concentration of the drug (7).

To place these results into context, the time to resistance for MIL was assessed in parallel, and was about 30 weeks (6 selection rounds), with a shift of the IC_50_ from 13 µM to 100 µM. We added for comparison, data from our previously published study with the same LdBPK_282 cl4 strain, in which we showed that the time to resistance to trivalent antimonials was 20 to 5 weeks (depending on the selection protocol) (7). Fig 1B shows clearly that the time to resistance was the longest for TCMDC-143345. The second evaluated parameter was the stability of the resistance phenotype. Both the WT line and Line D1 were maintained for 20 weeks without drug pressure and then challenged again with TCMDC-143345. From Fig. S2A, it can be observed that the withdrawal of the drug pressure for a prolonged period of time did not alter the susceptibility of line D1 to TCMDC-143345 (IC_50_ >25 µM). Thirdly, the susceptibility of intracellular amastigotes was assessed. TCMDC-143345 pressure was applied on THP-1 macrophages infected with line D1. Intracellular amastigotes of line D1 showed an IC_50_ of 30 µM versus 2 µM for the WT, confirming that the resistance selected at the promastigote stage was maintained at the amastigote stage (Fig. S2B). Fourthly, promastigotes of the resistant line D1, WT and D1-no drug (D1 line maintained for 20 weeks without drug pressure) were tested for their susceptibility to known antileishmanial compounds (MIL, Ampho B and Sb^III^) as well as one novel compound (Compound Y) that has a chemical structure related to TCMDC-143345. No cross-resistance was observed for Ampho B and Sb^III^ with similar IC_50_ values observed for all lines (Fig. S2C & E). There was increased susceptibility of the TCMDC-143345-resistant lines to MIL (Fig. S2D). Interestingly, all TCMDC-143345-resistant lines showed higher IC_50_ values for compound Y compared to the WT, thereby implying cross-resistance of the TCMDC-143345-resistant lines to compound Y (Fig. S2F). All IC_50_ values are shown in Data S1A. Finally, we looked at the *in vitro* fitness of the TCMDC-143345-resistant promastigotes in the absence of the drug. The resistant lines had a moderated but significant lower rate of growth than the wildtype (Fig. S2G and supplementary text): C was the most slowly growing line.

### Missense mutations in the gene encoding the dynamin-1 like protein LdoDLP1 can be found in independent selected resistant lines to TCMDC-143345

To identify genetic changes underlying the observed resistance to TCMDC-143345, we applied whole-genome sequencing (WGS) and we characterized genomic changes of nuclear DNA in the control WT and the resistant C and D1 lines at all steps of the selection process (see Fig. S1). First, we analyzed changes at the level of nucleotide sequence, *i*.*e*. single nucleotide polymorphisms (SNPs), and small insertion and deletions (INDELs). A total of 245 SNP variants not present at the start of drug selection were identified in all the lines (Data S1B). Among these, only 2 missense mutations with large and statistically significant changes in allele frequency were observed in line C (Fig. 2A & 2C). The first one appeared at 5 μM TCMDC-143345 with a gradual increase in the allele frequency from 0 to 1 in LdBPK_290029300, the gene encoding dynamin-1 like protein (LdoDLP1). This C:T missense mutation translates to a change from alanine at position 324 to threonine (Ala324Thr). The second significant change in the allele frequency in line C concerns another missense mutation G:T in chr6; the allele frequency shifts to 0.5 at 4 μM TCMDC-143345 in a conserved hypothetical protein (LdBPK_060013600) and stabilizes around 0.7 during further selection. Only one missense mutation with a significant change in the allele frequency has been observed in all D lines (Fig. 2B). It starts at 6 μM TCMDC-143345, changes from 0 to 1 and stays stable until the end of the selection pressure. Intriguingly, that mutation is also in the gene encoding LdoDLP1 (LdBPK_290029300), but results in a different change at the amino acid level compared to the LdoDLP1 mutation found in Line C; *i*.*e*., glutamate at position 655 changes to aspartate (Glu655Asp). These results were supported by using a testing approach inspired by burden tests for rare disease associations, confirming that the number of mutations in this gene is the most highly correlated with IC_50_ (see supplementary text and Fig. S3). No significant indels were detected in lines C or D(1-4) along the selection pressure.

**Figure 2.**
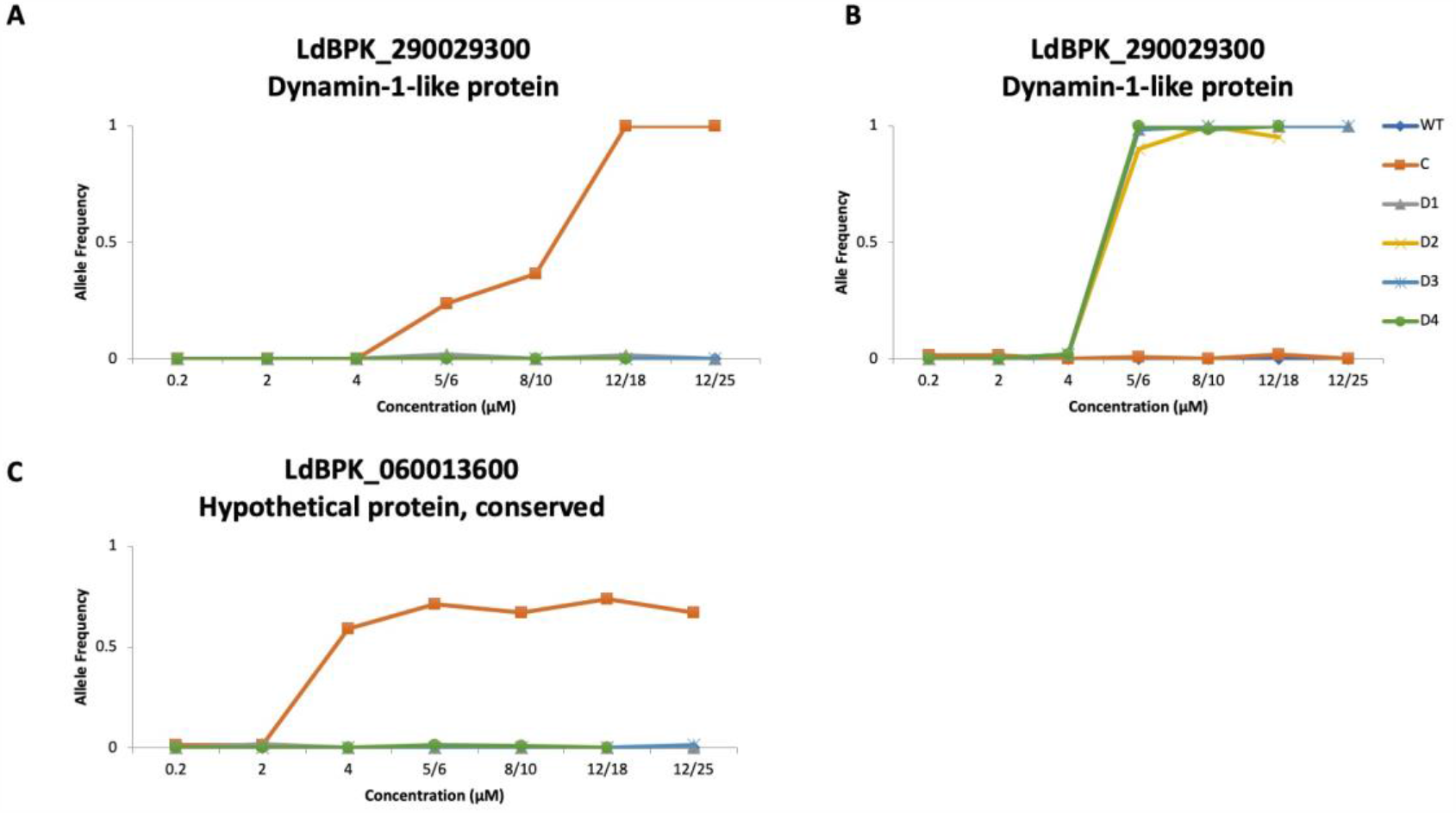
Major SNPs and their allele frequency change observed during selection of TCMDC-143345-resistance. (A) C:T missense mutation in position Ld29: 1000024 and (B) C:A missense mutation in position Ld29:999029C, both in the gene encoding dynamin1-like protein (LdoDLP1, LdBPK_290029300). (C) G:T missense mutation in position Ld06:353587 in a conserved hypothetical protein (LdBPK_060013600)

As we previously showed that aneuploidy and local copy number variations (CNVs) occur at an early stage of the selection process (8), we wondered whether it would be case in this experiment. Aneuploidy was already present before selection, but additional aneuploid chromosomes were observed around 4-6 µM TCMDC-143345 and final patterns of aneuploidy at the end of selection were rather different between Lines C and D1 (Fig. 3). Few CNVs were observed and the most striking were amplifications/deletions of large subtelomeric chromosomal stretches, also line-specific: in chr17 (D1) and chr30 (C) (Fig.S3B-C). More details on the aneuploidy and CNVs can be found in supplementary text.

**Fig 3.**
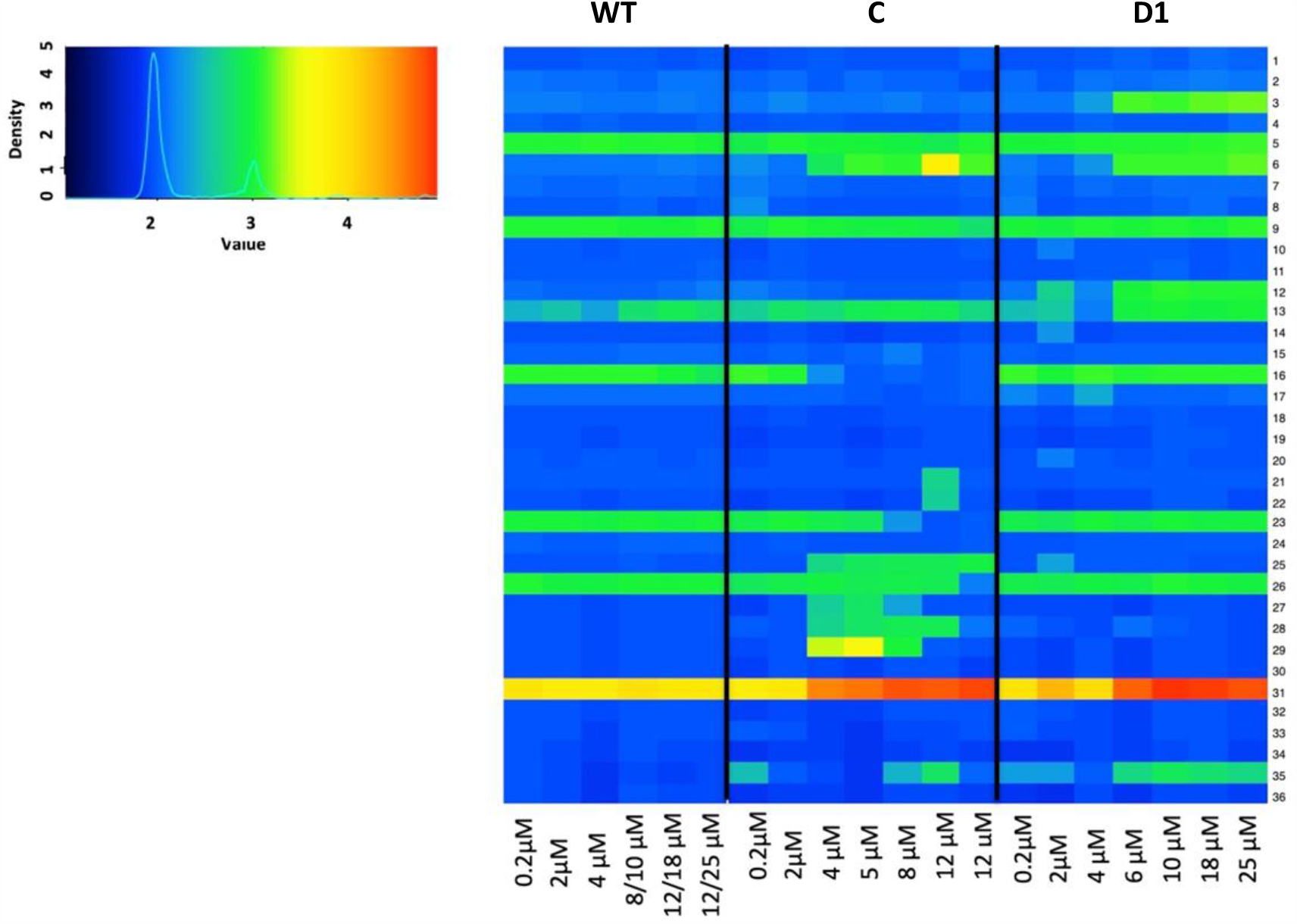
Somy changes in TCMDC-143345-resistant lines C and D1 and WT controls. Heat map representing the karyotype dynamics across the resistance selection of LdBPK_282 cl4 to TCMDC-143345. The color key shows the normalized chromosome read depth (S).

### The CRISPR-Cas9-mediated mutants and overexpression lines confirm the role of LdoDLP1 in the resistance to TCMDC-143345

As the only common missense mutation in the two independent resistant lines was found in LdoDLP1, we hypothesized that the genetic variation in this gene is responsible for the TCMDC-143345 resistance. To test this hypothesis, we recreated the identified mutations in wild-type promastigotes by means of a modified CRISPRS-Cas9 system described elsewhere (14) and we analyzed their resistance to TCMDC-143345. Detailed results of selection of transfected promastigotes and subsequent controls are shown in Supplementary text. Three clones were derived from each of the selected line (bearing the artificially introduced Ala324Thr or Glu655Asp mutations). The 6 clones were then submitted to a susceptibility test using a resazurin assay. The CRISPR-Cas9 engineered clones displayed a 4 to 5-fold increase in IC_50_ to TCMDC-143345 when compared to the WT LdBPK_282 cl4 or the uncloned parasites from control transfections #2 and #7 with a donor DNA lacking the missense mutations (DynWT1 and DynWT2 respectively - One-way ANOVA, P< 0.001), achieving an IC_50_ similar to Line C (Fig. 4A). Both the Ala324Thr (DynMut1 = mutation of line C) and the Glu655Asp (DynMut2 = mutation of line D) mutations had similar impacts on susceptibility to TCMDC-143345. Notably, no mutant clone displayed an IC_50_ similar to Line D.

**Fig 4.**
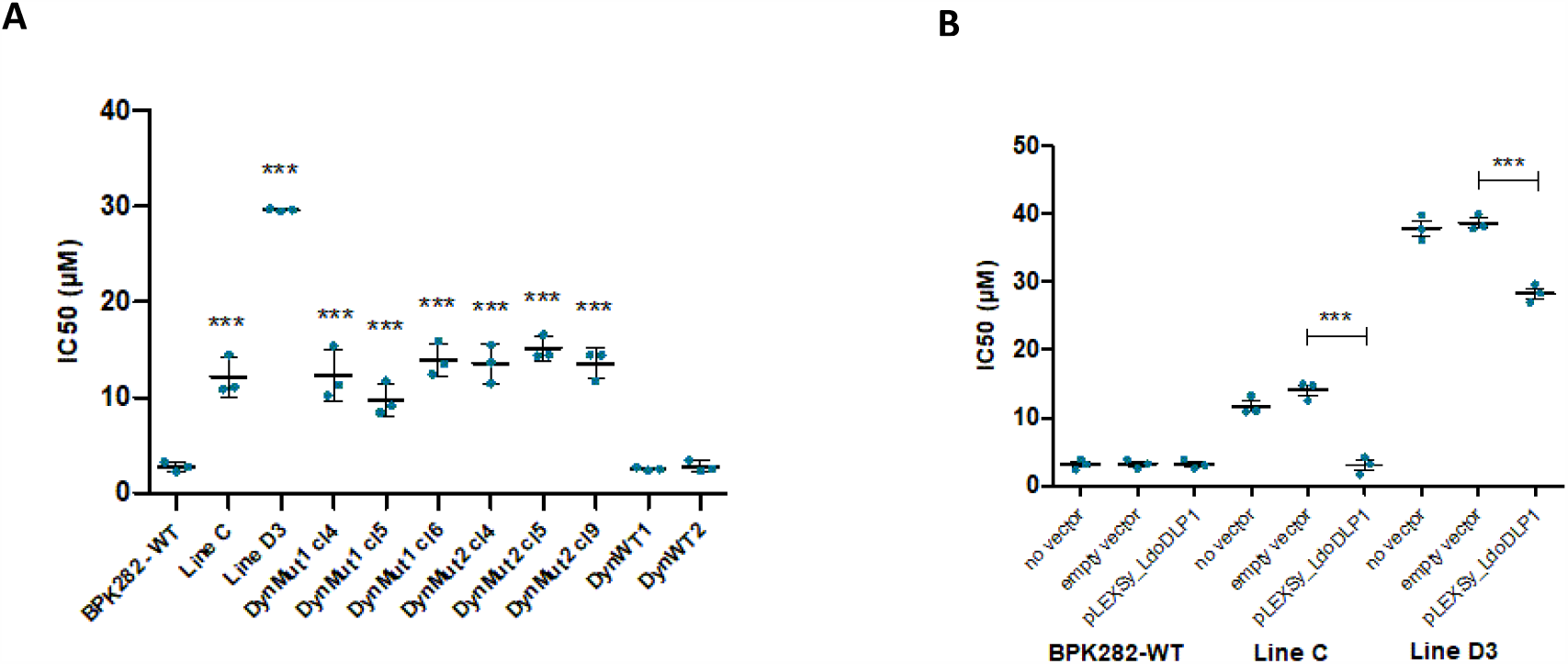
**A**. Susceptibility of 3 clones with the Ala324Thr (DynMut1 cl4, cl5 and cl6) or the Glu655Asp (DynMut2 cl4, cl5 and cl9) LdoDLP1 mutations introduced by CRISPR-Cas9. The WT LdBPK_282 cl4 and the unselected parasites transfected with DynWT1 or DynWT2 sgRNA and DNA templates were included as controls. **B**. Susceptibly of Lines C and D when overexpressing the WT LdoDLP1 gene. Lines represent the average and standard deviation of the three independent replicates (dots) of each experiment. *** = p ≤ 0.001 (ANOVA with Bonferroni’s Multiple Comparison Test).

In parallel to the CRISPR-Cas9-induced mutagenesis experiment, we also investigated the effect of over-expressing the wild type LdoDLP1 gene in lines C and D3. The transfection of an over-expression vector containing the wild type form of the gene completely abrogated resistance to TCMDC-143345 in Line C, reducing their IC_50_ to levels similar to the wild type LdBPK_282 cl4 (Fig. 4B - One-way ANOVA, P< 0.001). In line D3 however, while a significant reduction in IC_50_ was observed (One-way ANOVA, P< 0.001), parasites still demonstrated resistance to the compound, with an average IC_50_ of 29 µM. Altogether these results indicate that the mutations in the LdoDLP1 constitute the major genetic changes responsible for the observed resistance to TCMDC-143345 in both lines. Growth curves in the absence of drug pressure showed a similar growth rate between DynMut1 and DynMut2 and pT007 control, the WT line, which constitutively expresses Cas9 protein (Fig.S2H and supplementary text).

### Phylogenetic analysis suggests that LdoDLP1 plays a role in mitochondrial fission

LdoDLP1 belongs to the family of dynamin-related proteins (DRPs), a group of proteins that impacts the shapes of biological membranes by mediating fusion and fission events. Given the i) clear contributions of the Ala324Thr and Glu655Asp LdoDLP1 mutations to the TCMDC-143345 resistance phenotype and ii) the absence of detailed biochemical studies on leishmanial DRPs, a phylogenetic analysis was performed in order to learn more about the protein’s possible biological function. As was reported by Morgan and colleagues in their work on *Trypanosoma brucei* DLP1 (24), a sequence alignment followed by the construction of a rooted phylogenetic tree reveals that DRPs fall into different functional clades. Interestingly, LdoDLP1 clusters together with *T. brucei* DLP1 into the clade of DRPs involved in mitochondrial fission, (Figure S4), suggesting a role for LdoDLP1 in this biological process.

### TCMDC-143345 resistant lines exhibit lower mitochondrial membrane potential compared to susceptible, wild-type parasites

It has been well established that mitochondrial dynamics plays a role in the maintenance of normal mitochondrial membrane potential (MtMP) and cellular respiration (25)(26). Given the putative role of LdoDLP1 in mediating mitochondrial fission, we therefore evaluated whether the parasite lines containing LdoDLP1 mutations displayed changes in their mitochondrial activity. The MtMP and cell viability were simultaneously evaluated on logarithmic (day 2), early stationary (day 4) and late stationary promastigotes (day 7). In the presence or absence of TCMDC-143345 the overall trend was that cells with good viability in DR lines had decreased MtMP in comparison to the WT during the three days of testing (Two-way RM ANOVA, P< 0.001) (Fig 5A-B). However, the differences between WT and DR lines were bigger in logarithmic parasites (Fisher’s LSD test, P values < 0.0001) both in presence or absence of TCMDC-143345. Noteworthy, also on day2 the drug pressure dramatically alters the Mitotracker RFUs in the WT but barely in the case of resistant lines (see scatters plots Fig.5 A). We further evaluated if the CRISPR-engineered lines bearing the mutations Ala324Thr (DynMut1) and Glu655Asp (DynMut2) in LdoDLP1 had also diminished MtMP and it was confirmed for both mutations. Moreover, as with the *in vitro* selected DR lines the differences between LdoDLP mutant lines and the WT were larger during the logarithmic phase (Fig. S5). These results overall confirm that the resistant lines containing LdoDLP1 mutations have higher survival rate with the cost of moderate but significant decrease of the MtMP.

**Figure 5.**
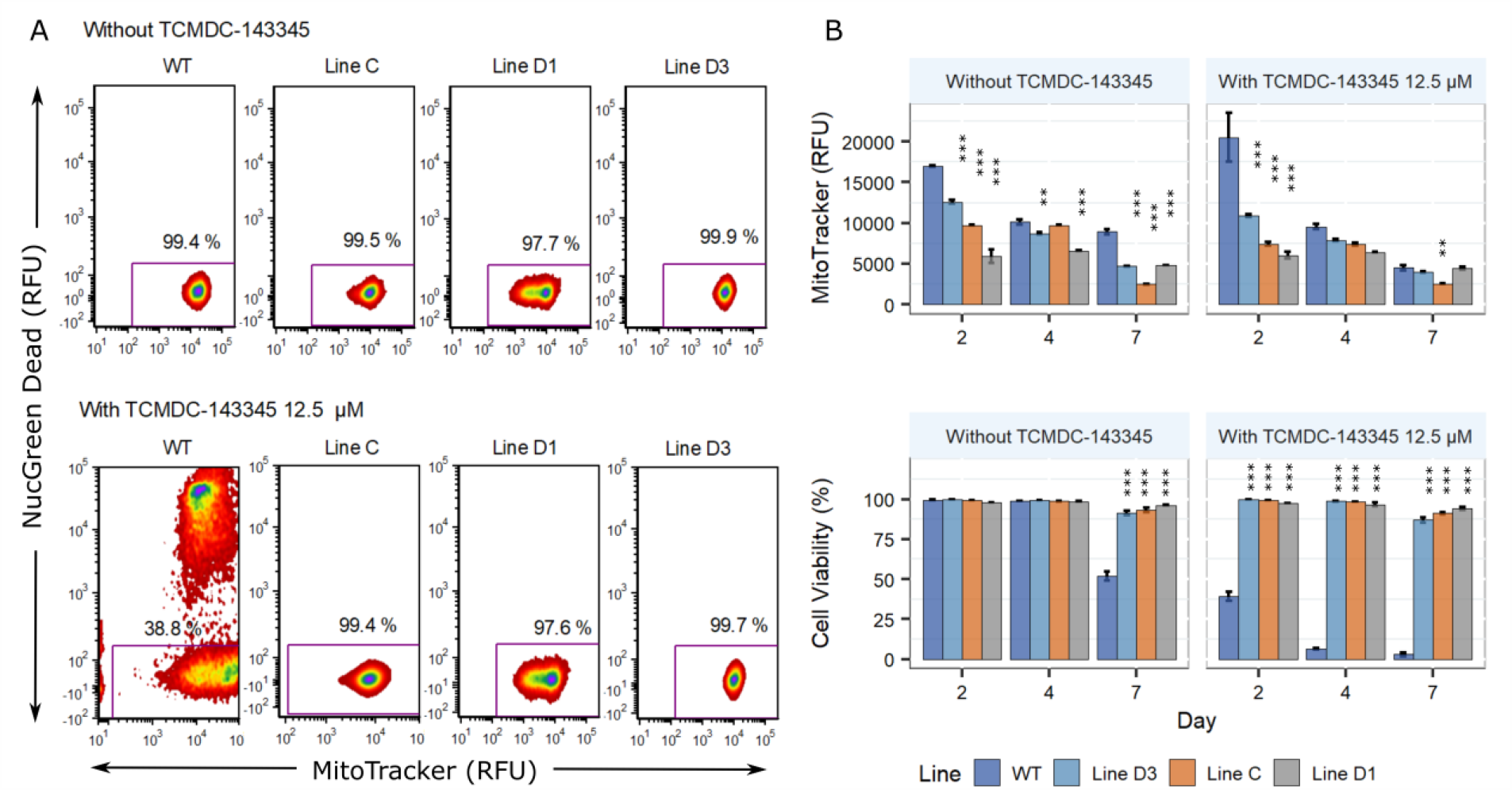
Mitochondrial membrane potential (MtMP) and cell viability of TCMDC-143345-resistant lines in promastigotes. (A) Density plots of the MtMP as measured by the fluorescence of the MitoTracker DeepRed and the percentage of cell viability as measured by the florescence of the NucGreen Dead. One plot of each line 2 days post treatment with 12.5 µM of TCMDC-143345 and their respective controls are shown. (B) MtMP in WT and resistant lines on days 2, 4 and 7 post treatment. All the resistant lines had diminished MtMP in the logarithmic phase of culture (Day 2) in comparison to the wildtype in presence or absence of TCMDC-143345. The differences between wild type and resistant lines were smaller in later days of incubation. Each bar represents the mean ± SEM of three biological replicates. Within each day, the asterisk in the top of the bars represent significant differences between the resistant line in comparison to the wild type (Fisher’s LSD test, P< 0.05).

### Homology modeling suggests a molecular basis for the putative impact of the Ala324Thr and Glu655Asp mutations on LdoDLP1 function

Given that the introduction of the LdoDLP1 Ala324Thr and Glu655Asp point mutations bestows TCMDC-143345 resistance onto the parasite and leads to an altered mitochondrial membrane potential, a structural model for LdoDLP1 was generated through homology modeling in an attempt to provide a molecular basis for these observations. LdoDLP1 contains all structural features characteristic of DRPs (Fig. 6, panel A): a neck domain (aka bundle-signaling element or BSE) consisting of three α-helices, a GTPase domain, an α-helical stalk domain that contains the dimerization interface, and a foot domain (aka ‘paddle’ or pleckstrin homology or PH domain). In contrast to various other DRPs, LdoDLP1 is devoid of the intrinsically disordered proline-rich domain (PRD).

**Figure 6.**
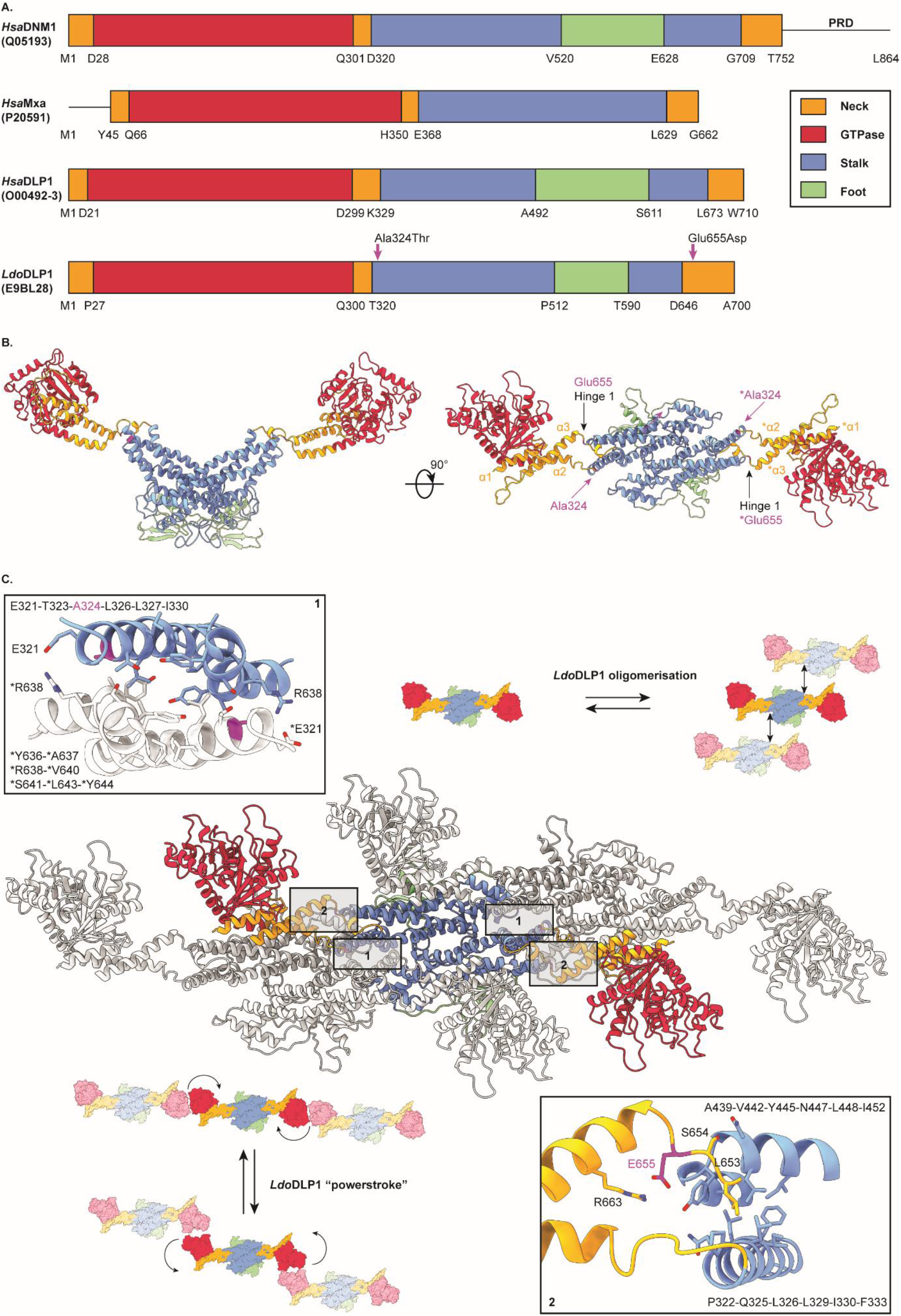
A homology model of LdoDLP1 suggests a molecular basis for the impact of TCMDC-143345 resistance mutations on protein function. (A.) Schematic representation of the primary sequences of *Homo sapiens* dynamin 1 (*Hsa*DNM1), *H. sapiens* myxovirus resistance protein 1 (*Hsa*MxA), *H. sapiens* dynamin-like protein 1 (*Hsa*DLP1) and LdoDLP1. The Uniprot identifiers are shown as well. The different domains are color-coded and their boundaries have been indicated. The LdoDLP1 Ala324Thr and Glu655Asp mutations associated with TCMDC-143345 resistance are indicated by the magenta arrows. (B.) Cartoon representation of the LdoDLP1 dimer homology model, color-coded as in panel A. The 3 α-helices of the neck domain, ‘Hinge 1’ and the positions of Ala324 and Glu655 have been indicated for convenience. ‘*’ is employed to indicate the second protomer of the LdoDLP1 dimer. (C.) The center displays a model of a higher order oligomer of the LdoDLP1. The central dimer is color-coded as in panel A, whereas the other two dimers are depicted in gray for reasons of clarity. The regions containing the Ala324Thr and Glu655Asp mutations are indicated by boxes ‘1’ and ‘2’, respectively. The insets provide a close-up view of the molecular interactions in these boxed regions. Relevant amino acids are shown in a stick representation. Ala324 (colored magenta) is proposed to be a part of the higher order oligomerization interface of LdoDLP1 (box 1), while Glu655 (colored magenta) is likely to play an important role in LdoDLP1’s “powerstroke” (box 2). ‘*’ is employed to indicate the second protomer of the LdoDLP1 dimer.

The Ala324Thr mutation maps onto a part of the stalk domain that is closely located to the α2-helix of the neck domain (Fig. 6B) and is known to be responsible for the higher-order oligomerization of DRP dimers, which is important for DRP function in fission events ((27) (28) (21) (29), more in-depth explanation in the Supplementary text). The Glu655Asp mutation is located in a region known as ‘Hinge 1’, which connects the stalk domain with the α3-helix of the neck domain (Fig. 6, panel B). ‘Hinge 1’ confers flexibility to DRPs, which is crucial for the so-called “hydrolysis-dependent powerstroke” underlying protein function ((29) (30) (31), more in-depth explanation in the Supplementary text). Hence, the LdoDLP1 mutations contributing to TCMDC-143345 resistance could i) alter the tendency of LdoDLP1 dimers to form higher-order oligomers (Ala324Thr) and ii) have a considerable impact on the flexibility of LdoDLP1’s ‘Hinge 1’ region (Glu655Asp), which would in turn be expected to affect protein function.

## Discussion

Humans and their pathogens are continuously locked in a molecular arms race and human interventions like chemotherapy have a further impact on parasites adaptations and counter-adaptations (1). This is well illustrated by the present study in which we (i) selected in *L. donovani* resistance to TCMDC-143345, a novel and potent anti-leishmanial compound that emerged from the Leishbox library of anti-leishmanial compounds (4) and (ii) characterized the extent of adaptations developed by the parasite.

Ten selection rounds (about 50 weeks) were necessary to obtain resistance to TCMDC-143345 in promastigotes, resistance was stable in the absence of the drug and it was also maintained in intracellular amastigotes. The same LdBPK_282 cl4 line was previously used for selecting and characterizing resistance to drugs used in clinical practice, hereby allowing comparisons about development of resistance to different compounds. Overall, it took more time to develop resistance *in vitro* to TCMDC-143345 (10 rounds, 50 weeks) than to MIL (present study, 6 rounds, 30 weeks) and to Sb^III^ (1-4 selection rounds or 5-20 weeks, depending on the protocol used)(7).

Whole genome sequencing is a powerful tool to identify molecular changes accompanying DR development in pathogens (7)(8, 32). In this study, in-depth genomic analysis revealed that TCMDC-143345 resistant lines were characterized by several aneuploidy changes, CNVs (see discussion in supplementary text) and SNPs. Especially the SNP analysis during the development of resistance turned out to be very informative. Out of 245 detected SNPs, only 2 missense mutations were fixed in 2 independent resistant lines and both are located in the gene encoding *L. donovani* dynamin-1 like protein (LdoDLP1). We used CRISPR-Cas9 to recreate the Ala324Thr or the Glu655Asp mutations in the LdoDLP1 gene of WT LdBPK_282 cl4 promastigotes. Hereby, we demonstrated that these mutations independently confer resistance to TCMDC-143345, leading to an IC_50_ in all mutant clones that is similar to Line C. Consistent with this, the over-expression of the WT LdoDLP1 gene completely abolished resistance to TCMDC-143345 in line C, while it had only a partial impact in the IC_50_ of line D3. These observations demonstrate that a loss-of-function mutations in LdoDLP1 gene are sufficient to provide a resistance to TCMDC-143345 in both lines. However, in line D this resistance is further increased by other uncharacterized modifiers acting in concert with the mutation in the LdoDLP1. Further work is needed to explore the functional impact of the specific aneuploidy changes, CNVs and SNPs observed in that line.

LdoDLP1 belongs to the family of dynamin-related proteins (DRPs) which are involved in several functions(24)(26): i) mitochondrial fusion (fusion of the inner mitochondrial membrane), ii) membrane dynamics of the outer chloroplast membrane, iii) endocytosis and vesicle trafficking in animals, iv) vesicle trafficking in plants, v) mitochondrial fission (fission of the outer mitochondrial membrane), and vi) plate formation and cell division in plants. DRPs cluster in different phylogenetic clades according to these functions(24) and our phylogenetic analysis showed a clear clustering of LdoDLP1 into the DRP clade involved in mitochondrial fission. The combined actions of mitochondrial fission and fusion govern mitochondrial dynamics, which underlie the organisation, copy number, form and function of mitochondria. The rates of these events are regulated according to the metabolic and/or developmental needs of the cell and in response to cellular stress or damage. While mitochondrial dynamics and cell division are not necessarily coupled in eukaryotes, the link between both processes seems to be very stringent in apicomplexan and kinetoplastid parasites. Both Apicomplexans and Kinetoplastids contain a single mitochondrion of which the fission is controlled by a single or a limited number of DLPs (33)(34). *Trypanosoma brucei* harbours two DLP paralogs (TbDLP1 and TbDLP2) (35), of which especially TbDLP1 seems to play a central role in linking the processes of mitochondrial fission, cytokinesis, and distribution of kinetoplastid DNA(24, 36–38). Interestingly, abrogation of TbDLP1 function in *T. brucei* blocks mitochondrial fission and cell division, again leading to parasite fatality(24, 36). Similar to other protozoan parasites, *Leishmania* spp. contain a single elongated mitochondrion and harbor a single DLP (24).

The clear clustering of LdoDLP1 with *T. brucei* DLP1 into the DRP clade involved in mitochondrial fission hinted towards a role for LdoDLP1 in mitochondrial dynamics, which is why we studied the mitochondrial membrane potential and cell viability of the TCMDC-143345 resistant parasite lines. In accordance with a proposed role of LdoDLP1 in mitochondrial fission, we found that all the TCMDC-143345-resistant lines (including the CRISPR-Cas9 generated LdoDLP1 Ala324Thr and Glu655Asp mutants) showed an altered mitochondrial activity. Interestingly, homology modeling suggests that the LdoDLP1 Ala324Thr and Glu655Asp mutations associated with TCMDC-143345 resistance are located in the protein’s oligomerisation interface and ‘Hinge 1’ region, respectively, two regions that are essential for general DRP (and thus LdoDLP1) function. Hence, these mutations are likely to influence protein function, which in turn might explain the observed impact on mitochondrial dynamics within the TCMDC-143345 resistant *L. donovani* parasites.

Whether LdoDLP1 is the molecular target of TCMDC-143345 or is part of a coping mechanism without being the direct target of TCMDC-143345 remains to be investigated. In this latter hypothesis, the LdoDLP1 mutations may have arisen to alleviate the drug’s detrimental effect on parasite viability. Within this context, it is interesting to note that leishmanial DLP was also proposed to be involved in the resistance profile of antimony- and miltefosine-resistant *Leishmania infantum* (39). In this proteomic study, *L. infantum* DLP was found to be down-regulated in the antimony- and miltefosine-resistant strains, although no information was gathered with regards to possible mutations in the protein. In the former hypothesis (*i*.*e*., LdoDLP1 is the molecular target of TCMDC-143345), the Ala324Thr and Glu655Asp mutations provide a direct escape to the drug’s deadly mode of action through LdoDLP1 binding. This hypothesis can equally be supported by the MtMP results. As of day 2 of drug exposure, we observed a drastic alteration in the Mitotracker RFUs for the WT parasite, whereas the Mitotracker RFUs remained unaffected for the resistant lines. This could be explained by the following scenarios: i) the LdoDLP1 mutations in the DR lines prevent the compound from binding LdoDLP1 (*i*.*e*., the mutations are located in the binding site for TCMDC-143345) or ii) they compensate for the effect that drug binding may have on LdoDLP1’s function in mitochondrial dynamics (TCMDC-143345 binds another LdoDLP1 ligand binding site). Clearly, in this hypothesis, the elucidation of the binding site and molecular interactions responsible for affinity and specific recognition between LdoDLP1 and TCMDC-143345 would shed relevant insights to be exploited in the design of new compounds with optimised potency. This would be especially interesting since DLPs from protozoan parasites are considered as drug targets because of their essentiality with regards to cell division and parasite growth (33).

Altogether, the results gathered in present study demonstrate the practical relevance of prospective DR studies. The time-to-resistance here shown for TCMDC-143345 is encouraging in the context of the shelf-life of that compound, but this should be complemented by *in vivo* studies. The demonstrated engagement of the unique leishmanial DLP in the resistance of *L. donovani* to TCMDC-143345 will allow the development of diagnostics targeting that gene to accompany further pre-clinical and clinical studies, if any and it will also guide further investigation on the mode of action. The absence of cross-resistance with other drugs currently used in clinical practice qualifies TCMDC-143345 for future combination therapy if the compound would reach that stage. It is still early to assess whether this mechanism of resistance is relevant against other chemical classes, yet the resistant strains selected become powerful tools to be employed with new chemical classes to ascertain whether they share biological space in terms of mode of action or resistance.

## Acknowledgements

This study has received funding from the European Union’s Horizon 2020 research and innovation programme under the Marie Sklodowska-Curie grant agreement N° 642609 and the Flemish Fund for Scientific Research (12Q8115N). MJ is supported by the Flemish Fund for Scientific Research (postdoctoral fellowship). GN is supported by the Flemish Ministry of Science and Innovation (SOFI Grant MADLEI). JAC, MB and MS are supported by the Wellcome Trust via their core funding of the Wellcome Trust Sanger Institute (grant 206194). The funders had no role in study design, data collection and interpretation, or the decision to submit the work for publication.

